# The Neurocognitive Mechanisms Underlying Narrative Schematic and Paraphrastic Transmission

**DOI:** 10.1101/2025.03.13.643115

**Authors:** Menghan Yang, Juan Zhang, Shijie Li, Chunming Lu, Lanfang Liu, Guosheng Ding

## Abstract

Narrative transmission serves multiple crucial functions, such as cultural preservation, knowledge accumulation, and consensus building, in human society. However, our understanding of its neurocognitive mechanisms remains limited. In this study, we combined a social transmission chain design with fMRI to investigate the dynamic changes of different narrative components occurring throughout transmission chains and to uncover the factors driving fidelity and distortion. A total of 58 participants were scanned as they listened to and subsequently recalled a story within a social transmission chain. We distinguished two types of transmission modes: schematic transmission, which prioritizes preserving the structural framework of the information, and paraphrastic transmission, which entails rephrasing the content while conveying its meaning. Behavioral results revealed a pattern in which paraphrastic transmission led to distortion and divergence, whereas schematic transmission remained more stable and convergent. Neural findings indicated that the transmission of story structure and content involved subsystems of the Default Mode Network (DMN) and specific subregions of the Hippocampus (HPC). While neural reinstatement between narrative listening and speaking failed to predict across-generation fidelity of content and structure, functional connectivity pattern similarity analysis showed that the DMN and HPC work as an episodic memory system supporting transmission performance. These findings highlight that narrative transmission is supported by distinct neurocognitive representations within episodic memory subsystems, which work collaboratively to sustain the transmission of narratives.

## Introduction

A narrative is a carefully structured depiction of events, experiences, or stories, frequently organized chronologically or in a cause-and-effect sequence. Due to their inherent ability to intriguingly and effectively convey information (Bruner, 1986; Oatley, 2016), narratives often serve as the transportation for the transmission of both individual and group knowledge and values (Hirst et al., 2018; Willems et al., 2020), and these social functions are further pivotal in human cultural accumulation and evolution (Acerbi et al., 2017; Kashima et al., 2019; Mesoudi, 2016; Wald, 2008).

Despite the irreplaceable social functions narrative transmission serves in preserving cultural heritage and fostering shared values, our understanding of the neurocognitive mechanisms underlying this process remains limited. It has been widely believed that human episodic memory plays a crucial role in narrative transmission, allowing individuals to retain and recall stories, and further preserving, passing down, and maintaining a shared understanding (Hirst et al., 2018). However, episodic memory is inherently reconstructive (Bartlett, 1932; Foudil et al., 2020; Marsh, 2007), leading to distortion, misinformation, or biased transmission (Coronel et al., 2020; Mesoudi et al., 2006; Willard et al., 2016). This raises critical questions about the factors driving both the distortion and preservation of information during narrative transmission. Specifically, the ways in which the reconstructive nature of episodic memory affects transmission fidelity, and to what extent, remain unknown.

Recently, increasing evidence shows that humans have distinctive representations of different components of episodic memory, such as schema and details (Heusser et al., 2021; Lynn & Bassett, 2020; Robin & Moscovitch, 2017). For example, narrative schema serves as a scaffold in encoding and retrieving episodic memories, supporting the overall cognitive process of narrative structure (Alaoui-Soce & Tamir, 2023; Baldassano et al., 2018; Masís-Obando et al., 2022; Reagh & Ranganath, 2023), while details contribute to constructing individuals’ situation models, enhancing personal understanding of narrative plots, and elaborating story content (Marsh, 2007; Robin & Moscovitch, 2017; Zwaan & Radvansky, 1998).

Based on previous evidence, here, we propose that narrative transmission has two types of transmission modes: **schematic transmission** and **paraphrastic transmission**. Each mode is underlined by distinctive human representations in episodic memory components, and plays different functions in information retrieval and narrative transmission. Schematic transmission aims to convey the core or essential framework of a story, organizing elements in the narratives while leaving out most of the finer details. It focuses more on conveying the abstract structure of a story rather than providing a verbatim or detailed reproduction of the entire narrative, therefore allowing for a more efficient and concise transmission of information. Paraphrastic transmission underlines the retrieval of narrative content or plots in a rephrased form. It normally involves restating the narrative in different words, often with the effect of simplifying complex language or concepts based on individuals’ interpretations, making the story more accessible. As a support to the existence of the two type modes, Heusser et al. (2021) applied a geometrical method to distinguish narrative structure and content, and found that narrative recall is a structure-based yet details-deviated process. Meanwhile, evidence also suggests that stories with familiar structures hold an evolutionary advantage over those with unfamiliar formats (Alaoui-Soce & Tamir, 2023). Nevertheless, it remains uncertain how these two mechanisms collaboratively or uniquely influence the transmission of narratives among multiple individuals and, in the end, shape memory outcomes within a group.

Neural research offers compelling evidence for the distinct representations of narrative structure and content. Existing findings suggest that the hippocampus serves as a hub for episodic memory, where various features of episodic memory converge (Backus et al., 2016; Cohn-Sheehy et al., 2021; Milivojevic et al., 2016). The neural representations for detailed, gist, and schematic information within episodic memory are spatially segregated (Masís-Obando et al., 2022; Robin & Moscovitch, 2017), and associated with certain functional connectivity between the hippocampus and neocortex (Arnold et al., 2018; Cooper & Ritchey, 2019; Reagh & Ranganath, 2023). Specifically, the activity in the posterior hippocampus and neocortex is related to detailed and specific information representation, and the representation of schematic information predominantly occurs in the ventromedial prefrontal cortex (vmPFC). Meanwhile, recently, researchers have divided the default mode network into different subsystems (Schaefer et al., 2018), and these subsystems play distinct roles in episodic memory encoding and retrieval (Ritchey & Cooper, 2020). Together, converging evidence has stressed the separating roles of the hippocampus and the default mode network in the representation, encoding, and retrieval of different components of episodic memory. However, due to the complexity of narratives and the challenges of neural data acquisition, most research has focused on narrative encoding and retrieval within individuals (Zadbood et al., 2017). In fact, compared to intrapersonal transmission, social transmission relies on the exchange of information via memory, imitation, and communication, and therefore has variable fidelity and may become distorted, simplified, or enhanced during interpersonal transmission (Almaatouq et al., 2020; Legare, 2017)(Freire & Verschure, 2024; Montrey & Shultz, 2020; Saldana et al., 2019) (Freire & Verschure, 2024; Montrey & Shultz, 2020; Saldana et al., 2019). While prior studies have illuminated how individual episodic memory functions, how these internal mechanisms shape the transmission of information across individuals remains unclear. To address this gap, our study examines whether the neural processes underlying episodic memory support high-fidelity narrative transmission or contribute to distortions as narratives propagate across multiple generations.

To answer the above questions, we applied the **Transmission Chain Paradigm** (Bartlett, 1932; Kirby et al., 2008; Miton & Charbonneau, 2018) within a functional Magnetic Resonance Imaging (fMRI) framework to systematically examine how information is transmitted along a linear sequence of individuals and to uncover the underlying neurocognitive mechanisms. This paradigm simulates the real-world propagation of narrative, music, social, and cultural information, akin to the classic “Telephone” game (Breithaupt et al., 2022; Coronel et al., 2020; Lumaca et al., 2019). In this setup, participants listen to a narrative, recall it, and then relay their recollection to the next participant in the chain. This iterative process allows for a structured comparison of narrative fidelity across multiple generations. By utilizing this paradigm, we aimed to investigate how content and structure are preserved or distorted during the transmission process.

We recruited 58 participants, forming two transmission chains with four generations in each chain (for details, please refer to the Method section). In the scanner, each participant listened to the story generated by one of the preceding participants in the last generation and subsequently retold the story (see Figure 1A). To investigate how two representational modes—schema representation and content representation—shape narrative transmission, we compared the generational distortion patterns of **story structure** and **content** throughout the transmission process. Furthermore, we analyzed neural patterns to explore how the human brain supports and drives the propagation of stories through word-of-mouth across individuals.

**Figure 1.**
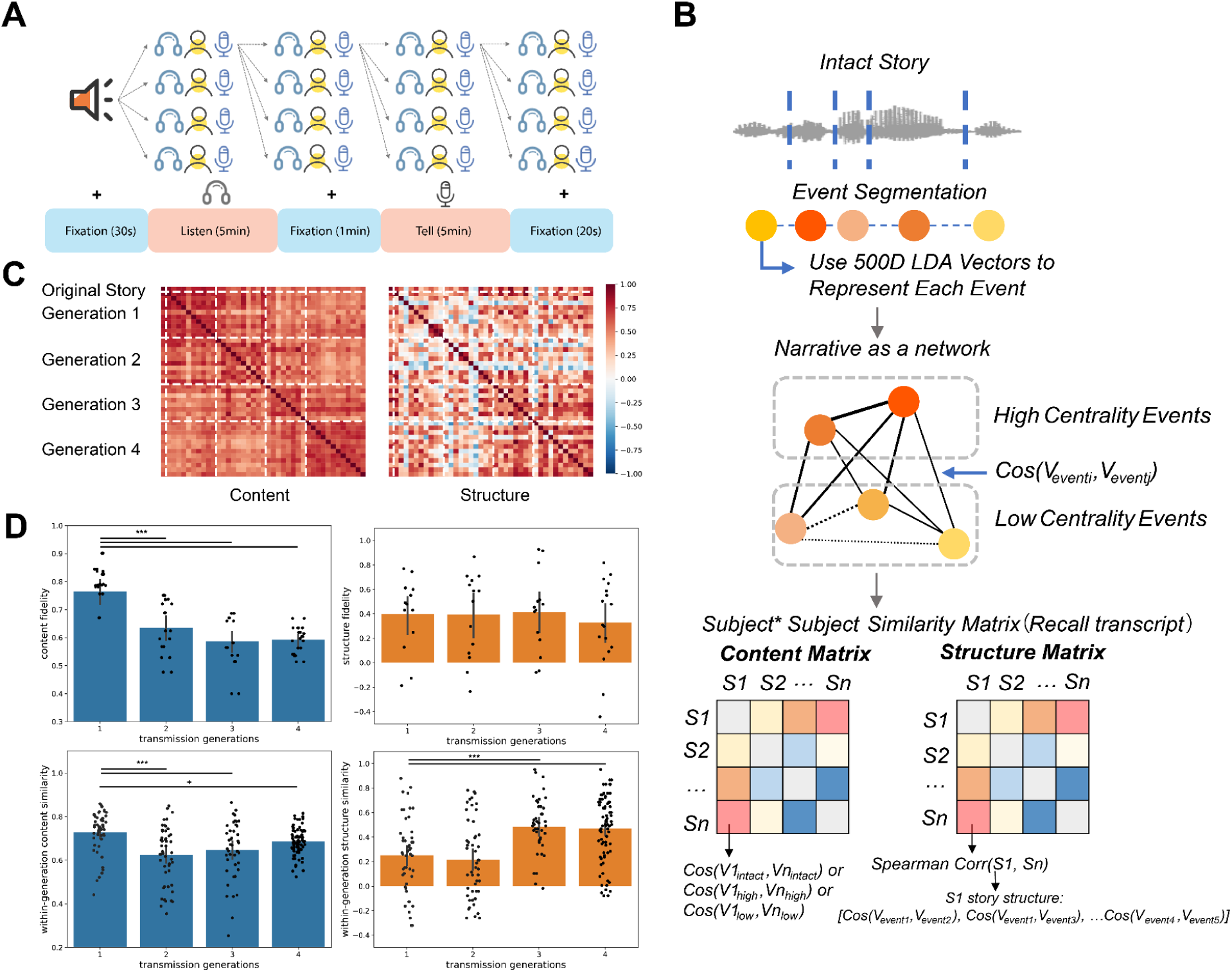
Schematic illustrations of the experimental paradigm and methodology, and behavioral narrative transmission results. (A) Transmission chain paradigm and individual experimental procedure in fMRI scanner. In the scanner, each participant listened to a story produced by a preceding participant from the previous generation and then recounted it. (B) Using Event Segmentation and LDA language model to decompose narrative content and structure and distinguish high- and low-centrality events. Content was represented as vectors of intact stories, whereas the structure of each story was captured through vectors of all the edges within the narrative. (C) Narrative paraphrastic (content, left) and schematic (structure, right) transmission across generations. Distinct patterns in the transmission of narrative content and structure were identified. (D) Changing patterns of Transmission fidelity (retold stories compared to the original story) across generations and within-generation similarity. We showed a pattern of distorted and divergent paraphrastic transmission of narrative content, alongside a convergent and stable transmission of narrative structure. +: 0.01 < *p* < 0.05; ***: *p* < 0.001, generation comparisons were under Bonferroni correction.

## Results

### The Distortions and Fidelity of Narrative Structure and Content

We first asked how various components of a story were distorted as participants passed the narrative through transmission chains. We utilized the computational language model and network analysis to answer this question. According to the *Event Segment Theory* (Kurby & Zacks, 2008; Zacks, 2020) and following the method of Lee & Chen (2022), the original story was divided into 5 events. Subsequently, the narratives recalled by each participant were also segmented into five events to align with the original storyline. A computational language model (Latent Dirichlet Allocation, LDA, see Method for more details) was used to vectorize each event. Each narrative (both original and recalled ones) can be represented as a narrative network, quantified by measuring the cosine similarity between event vectors. To assess content similarity, we computed the cosine similarity among the **content vectors of intact stories**. The structure of each story was represented by **the vector of all the edges** within a narrative network. Structure similarity was quantified by using the Spearman correlation between the **structure vector representations (network’s edges) of the stories** (Figure 1B).

We first observed overall distinctive changing patterns of narrative content and structure transmission in both chains by plotting content and structure transmission similarity matrices (Fig.1C and S1A). Previous studies indicated that specific details are more susceptible to distortion than schematic information (Robin & Moscovitch, 2017), therefore, we investigated whether the **narrative fidelity** changes across the transmission chains. We defined both content and structure fidelity by calculating the similarity between the original and recalled narratives. Our findings indicated that, after controlling for variations in transmission chains, content fidelity significantly declined (*F*_(3, 53)_ = 18.189, *p* < 0.001). In contrast, structural fidelity did not exhibit significant generational differences across transmission chains (*F*_(3, 53)_ = 0.252, *p*s > 0.01) (see Fig. 1D upper). The effects observed across different chains are illustrated in Figure S1B.

Specifically, we asked whether different events in the narratives exhibit varying degrees of fidelity, as previous research showed that items with higher centrality can be retrieved more easily (Coman et al., 2016; Lee & Chen, 2022). Therefore, further analysis is aimed at exploring the transmission patterns of different types of events in the narratives. To investigate this question, we adopted the methodology from previous research (Lee & Chen, 2022), identifying event nodes in the original narrative with higher centrality as high-centrality events, characterized by greater semantic similarity with other events in the story. Events with lower centrality were classified as low-centrality events. Consistent with previous findings, our results also revealed that high centrality events showed overall higher similarity in all 4 generations in both chains than low centrality events (*p* < 0.001, see Figure S2).

Lastly, we examined whether narrative content and structure **converged** within one generation as transmission progressed along the chains, which could be the mechanisms of human collective memory (Chang et al., 2024; Coman et al., 2016; Hirst et al., 2018). We, therefore compared within-generation content and structure similarity across different generations ( Fig. 1D lower). The results revealed an increase in within-generation structural similarity (*F*_(3, 213)_ = 13.560, *p* < 0.001) during transmission, whereas content similarity exhibited significant generational differences (*F*_(3, 213)_ = 9.980, *p* < 0.001) —specifically, comparisons of subsequent generations to the first generation showed a decline in within-generation content similarity (*β*s < -0.047, *p*s < 0.079). Detailed chain differences are exhibited in Figure S1C. Collectively, these findings suggest a pattern of distorted and divergent narrative paraphrastic transmission, along with a convergent narrative schematic transmission.

### Neural Representations of Narrative Structure and Content during Inter-Person Transmission

Despite the distinct behavioral patterns observed between narrative paraphrastic and schematic transmission, it is still unknown how our brain supports this process. To further answer this question, we used intersubject representational similarity analysis (IS-RSA, Figure 2A) by computing the correlations between subject-by-subject neural and text representation dissimilarity matrices (RDMs), to explore the shared and distinctive neural representations of structure and content transmission.

**Figure 2.**
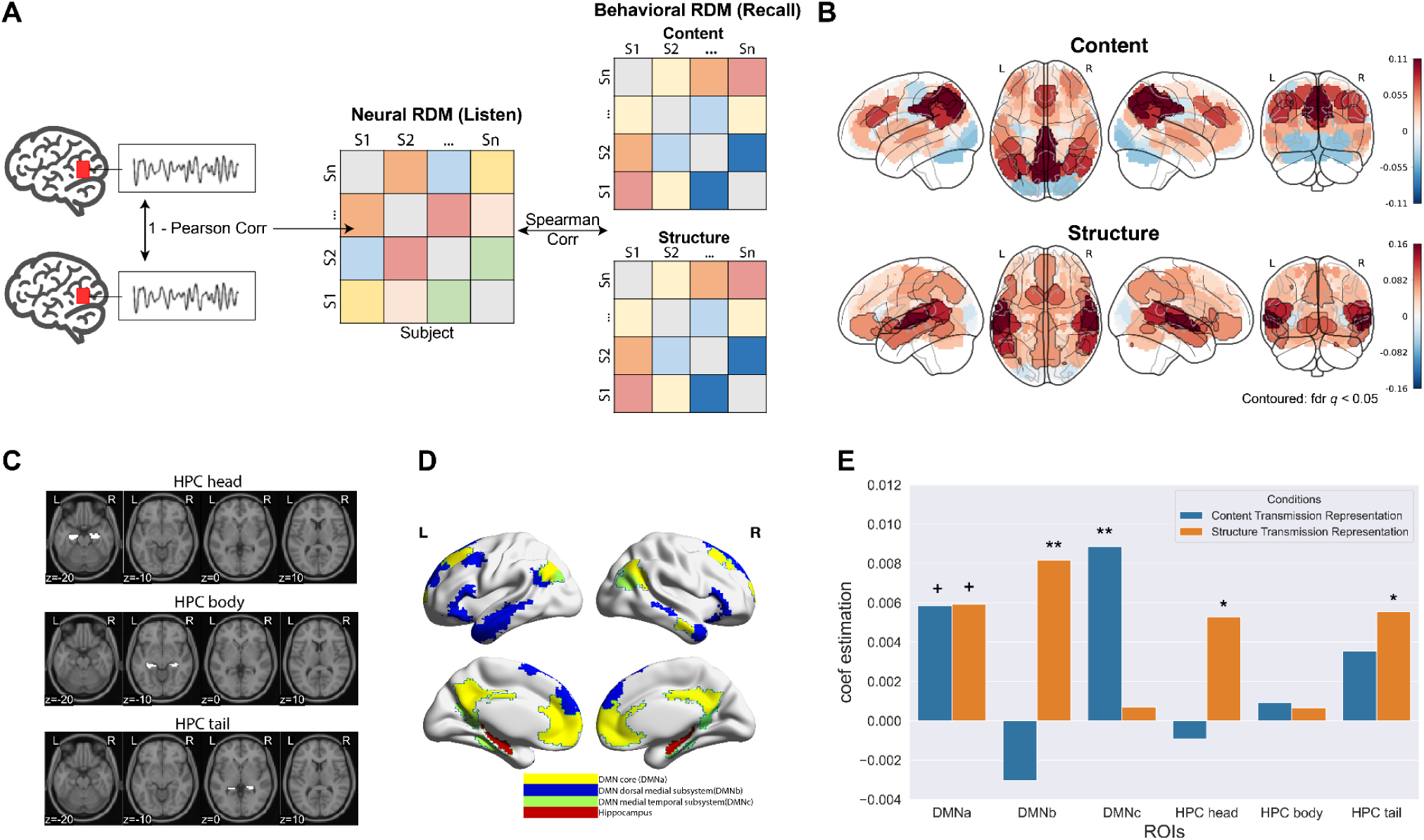
Neural representations of narrative structure and content during inter-person transmission. (A) Schematic illustration of Intersubject Representational Similarity Analysis (IS-RSA). (B) Neural representations of narrative structure and content across the whole brain. DMNa demonstrated overlapping representations for both content and structure. Content transmission exhibited stronger and more widespread representation in the superior lateral occipital cortex and parietal regions, whereas structure transmission was primarily localized in the temporal lobe, with both representations overlapping in the precuneus. The results were FDR-corrected for multiple comparisons across 50 ROIs. (C and D) DMN subnetworks and HPC subregions. (E) Neural representations on HPC subregions and DMN subnetworks. DMNb and the anterior HPC were specifically linked to content transmission, while DMNc and the posterior HPC were distinctly associated with structure transmission.

The global results indicated that both content and structure transmission were involved with Default Mode Network-related regions, and specifically overlapped at the precuneus (Figure 2B). Meanwhile, content-transmission representation showed a stronger and more extensive representation in the Superior Lateral Occipital Cortex and Parietal areas, while structure transmission representation occurs in the temporal lobe. We also observed that there existed content and structure representations in the prefrontal cortex, and structure-related representation was stronger in the ventromedial part while content-related one was in the dorsomedial subregions.

Previous studies suggest that different subregions of the hippocampus (anterior and posterior) have functional connections with the medial prefrontal cortex and posterior medial cortex, which are related to gist and detail processing in episodic memory (Robin & Moscovitch, 2017), implying separate functions within subareas of the hippocampus and DMN. Meanwhile, evidence also showed that these core brain regions within the default network were involved in encoding different types of information in episodic memory, such as scene schemas, protagonist information, and contextual differences (Reagh & Ranganath, 2023). We then specifically examined how episodic memory subnetworks and subregions represented content and structure transmission. Subregions of the Hippocampus (head, body, and tail, Figure 2C) and Different DMN subsystems (Core, Dorsal Medial, and Medial Temporal Subsystem, Figure 2D) were chosen as the regions of interest (ROI). We observed shared representations in DMNa for both content and structure, while distinctive representations were shown in DMNb (content transmission related) and DMNc (structure transmission related). Meanwhile, the anterior and posterior hippocampus also showed content and structure representations respectively (Figure 2E). Together, these findings highlight the distinct roles of DMN subsystems and hippocampal subregions in processing different narrative elements, suggesting that these neural mechanisms may facilitate narrative transmission across generations.

### HPC and DMNs Support High Fidelity Transmission As an Episodic Memory System

While our above results indicate the distinctive roles of HPC and DMNs in narrative representation, it is unclear how these regions and networks support the transmission of narratives across generations. Previous evidence indicated neural reinstatement works as a potential mechanism to support episodic memory encoding and retrieval and predict memory performance (Bird et al., 2015; Cohn-Sheehy et al., 2021; Masís-Obando et al., 2022; Nau et al., 2024), we then investigated whether neural reinstatement between/within individual subregions and subnetworks could predict the fidelity of content and structure transmission. We tested whether (1) temporal neural reinstatement between individual listening and retelling within the same region/subnetwork, (2) temporal neural reinstatement between listening and retelling across regions and subnetworks, or (3) temporal neural similarity within each region or subnetwork across generations (first-generation listening compared to other generations’ listening and speaking) could predict narrative fidelity. However, neither content nor structure fidelity could be significantly predicted by these neural reinstatement results (Figure S3, see supplementary information).

Despite the minor effects within each single brain region and subnetwork on across-generation fidelity prediction, there still existed the possibility that the functional connectivity patterns could serve as a fidelity predictor across generations. Under this analysis, we hypothesized that HPC subregions and DMN subnetworks work as an episodic memory system (Cooper & Ritchey, 2019; Ritchey et al., 2014; Ritchey & Cooper, 2020), and therefore, the similarity between their connectivity patterns and first-generation connectivity patterns could support narrative transmission.

To validate the above hypothesis, we separately computed functional connectivity between the hippocampus head/body/tail and default mode subnetwork during both the listening and speaking phases for each participant. We then calculated the similarity of FC patterns between each participant and the averaged FC patterns in the first generation, and examined whether this FC pattern similarity could predict high-fidelity transmission of content and structure across different generations (Figure 3A).

**Figure 3.**
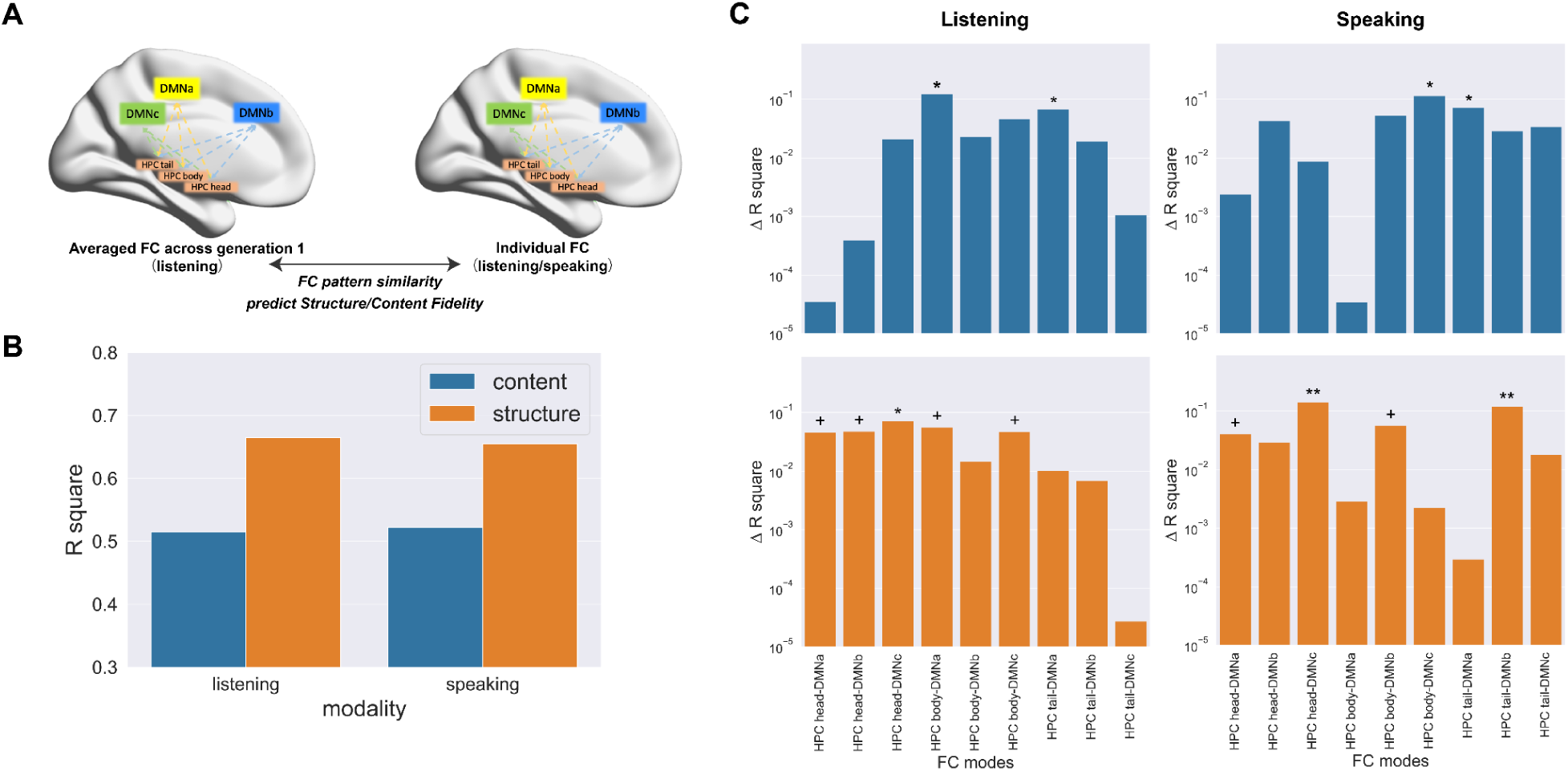
HPC and DMNs Support High Fidelity Transmission As an Episodic Memory System. (A) Illustration of how FC pattern similarity could predict fidelity. (B) The overall percentage of variance explained by FC similarity patterns in content/structure regression models (R^2^) in different modalities. (C) Prediction of different FC modes on fidelity during listening/speaking. The Y-axis represents the variance removed by each individual FC mode when excluded from the model. The Y-axis values were scaled using a logarithmic transformation for visualization purposes. The subregions of the HPC and DMN subnetworks function as a unified episodic memory system rather than operating independently to support the fidelity of both content and structure.

The results showed that during the phase of story listening, FC pattern similarity could predict the fidelity of structure (*β* = 0.375, *p* = 0.022). Although the effect on content fidelity is not significant (*β* = 0.244, *p* = 0.146), it became stronger (*β* = 0.255, *p* = 0.086) after controlling for the chain differences, with a similarly stronger effect on structure fidelity (*β* = 0.364, *p* = 0.012). In the meantime, we did not observe any significant results during the phase of story retelling for either content (*β* = 0.165, *p* = 0.176) or structure (*β* = 0.169, *p* = 0.789).

Furthermore, we investigated how different types of FC pattern similarity could jointly and individually contribute to high-fidelity content and structure transmission. We used a fixed-effects regression model to separate the effect of different FC pattern similarities. We observed an overall high model fit (R^2^_content model_ = 0.515, R^2^_structure model_ = 0.665, Figure 3B left) in both content and structure fidelity prediction. We further removed each FC individually from the model to test the variance explained by different FC modes. We observed the involvement of DMNa connected to the HPC body and tail in predicting content fidelity (*p*s < 0.069), and the roles of the HPC head and body connected to multiple subnetworks in predicting structure fidelity (*p*s < 0.073, Figure 3C upper and lower left).

We repeated this analysis on the speaking data (comparing FC connectivity modes when participants retold the stories to the FC connectivity mode when the first-generation participants were listening), and still observed an overall high model fit (R^2^_content model_ = 0.522, R^2^_structure model_ = 0.655, Figure 3B right). After removing each FC from the model, we found that the HPC body connected to DMNc, as well as the connection between the HPC tail and DMNa, significantly explained the variance in across-generation content fidelity prediction (*p*s < 0.059). Meanwhile, the involvement of DMNb (connected to the HPC tail and body) and the HPC head (connected to DMNa and DMNc) predicted across-generation structure fidelity (*p*s < 0.093; Figure 3C, upper and lower right).

Together, combining the above results, we demonstrated that the subregions of HPC and DMN subnetworks work as an episodic memory system to support content and structure fidelity.

## Discussion

Here we applied the narrative transmission chain and neuroimaging to explore the neurocognitive mechanisms of interpersonal narrative transmission. Our behavioral results demonstrate a pattern of distortion and divergence in narrative paraphrastic transmission, with convergence in narrative schematic transmission during oral storytelling. Specifically, while the fidelity of story content is relatively low and subject to continuous divergence across generations, high-centrality events tend to maintain stability, in contrast to the greater variability observed in low-centralized events. In contrast, the story structure demonstrates stable fidelity and higher within-generation similarity across the transmission. Furthermore, our neural results revealed that DMN subsystems and hippocampal (HPC) subregions play distinct roles in representing content and structure during narrative transmission, with overlapping activity in the precuneus. Specifically, DMNa exhibited shared representations for both content and structure, whereas DMNb and the anterior HPC were uniquely associated with content transmission, while DMNc and the posterior HPC were selectively involved in structure transmission. Finally, the HPC subregions and DMN subnetworks work cooperatively as an episodic memory system, due to the crucial role FC patterns play in predicting the fidelity of both content and structure across transmission generations. Collectively, these results suggest that a structured transmission model centered around high-centrality events emerges, even as specific details undergo modification, during which the different DMN subsystems and hippocampal subregions engage in both collaborative and independent functions of the content and structure representation and transmission.

Our behavioral results are in line with prior research, where studies employing word-embedding spaces to model participants’ oral recall of the same narrative demonstrated that recollections of details were notably coarse, albeit retaining a general sense of the “spatial attributes” or trajectory of the story progression (Heusser et al., 2021). This study further expanded research results from an “interpersonal” perspective, indicating that the structure and content of narratives reflect varied cognitive processes, suggesting that individuals likely employ distinct cognitive strategies—namely “**graph learning**” and “**situation construction**”—to accomplish narrative schematic and paraphrastic transmission (Costabile, 2016; Lynn & Bassett, 2020; Masís-Obando et al., 2022; Zwaan & Radvansky, 1998). The perception, encoding, and transmission of story structure appear to hinge on social learning, wherein individuals imitate and optimize structural elements for effective and efficient transmission. This learning modality is not only pivotal in acquiring structured knowledge (Lynn & Bassett, 2020) but also manifests in the schema of stories and social information (Augoustinos & Innes, 1990; Orianne & Eustache, 2023; Wertsch, 2004). Conversely, the understanding and transmission of narrative content rely more profoundly on sensory representations (such as visual reactivation) and imaginative engagement, making certain details more susceptible to distortion (Geib et al., 2017; Zadbood et al., 2017).

Furthermore, our segregated neural representation findings provided further evidence for this hypothesis. Existing research indicates that the hippocampus may serve as a central hub for episodic memory, where various features of contextual memory converge (Backus et al., 2016). Studies have shown that the representational similarity of the hippocampus increases when processing coherent narratives, suggesting the hippocampus plays a vital role in narrative situation construction (Cohn-Sheehy et al., 2021). Furthermore, research has indicated that within episodic memory, the brain spatially separates representations of detailed information, thematic content, and schematic information (Robin & Moscovitch, 2017). Specifically, the posterior hippocampus and neocortex influence the representation of detailed and specific information, while the anterior hippocampus is involved in representing gist information (Robin & Moscovitch, 2017). In contrast, the representation of schematic information predominantly occurs in the medial prefrontal cortex (Baldassano et al., 2018; Gagnepain et al., 2020; Masís-Obando et al., 2022; Reagh & Ranganath, 2023). Further, Kauttonen et al. (2018) employed multivoxel pattern analysis to identify activation patterns in higher-order cortical regions—including the precuneus, angular gyrus, cingulate gyrus, and frontal cortex—when representing essential scenes from films, supporting the roles these regions play in providing context and background during contextual memory retrieval. Our neural evidence suggests that the DMN core system (including posterior cingulate and anterior medial prefrontal cortex) showed the neural representations of both content and structure during narrative inter-person transmission. Meanwhile, anterior and posterior HPC and DMNb (Dorsal Medial Subsystem, including dMPFC, temporoparietal junction, lateral temporal cortex, etc.) were involved in content representation, while DMNc (Medial Temporal Subsystem, including vmPFC, posterior inferior parietal lobule, and parahippocampal cortex, etc.) showed the neural representation of schematic transmission. These findings on segregated DMN spatial representations validate the distinct functions of default network subsystems, highlighting the diverse neural mechanisms underlying schematic and paraphrastic transmission.

Although this study did not find patterns of reinstatement or temporal similarity within the same individuals that could predict story transmission fidelity, it underscores the importance of hippocampal connectivity with the neocortex in collaboratively supporting the recollection and transmission of semantic and schematic information. Previous research has highlighted the importance of neural reinstatement in successful memory retrieval (Baldassano et al., 2017; Bird et al., 2015; Cohn-Sheehy et al., 2021; Masís-Obando et al., 2022; Nau et al., 2024; Ritchey et al., 2013). However, factors such as interpersonal transmission in our study may limit its effectiveness, making neural reinstatement within the same individual and region an unreliable predictor of successful transmission. Therefore, we explored whether functional connectivity patterns could serve as an alternative predictor, under the assumption that episodic memory subnetworks operate as a dynamic system (Cooper & Ritchey, 2019; Ritchey & Cooper, 2020). Indeed, the distinct representational patterns of memory features are linked to the functional connectivity between the hippocampus and neocortex (Arnold et al., 2018; Cooper & Ritchey, 2019). Previous research also highlighted the importance of hippocampal-neocortical connections in contextual memory retrieval (Ren et al., 2018). This evidence, together, underscores the role of hippocampal-neocortical interactions in effective memory retrieval. Given that functional connectivity patterns predict various cognitive and behavioral traits (Meskaldji et al., 2016; Rosenberg et al., 2016) and their similarity has been linked to interpersonal closeness (Hyon et al., 2020), we employed functional connectivity similarity analysis to further investigate hippocampal-neocortical interactions in memory processes. Our findings demonstrate that FC pattern similarity supports both content and structure transmission during encoding and retrieval, challenging the idea that episodic memory fidelity relies solely on neural reinstatement and underscoring the complexity of memory processes involved in interpersonal storytelling.

In general, our work adopts an intergenerational transmission paradigm within the framework of social representation, extending experimental investigations from behavioral analyses to neural mechanisms. This shift facilitates a deeper understanding of representation differences and shared cognitive processes of social information transmission at the neurobiological level. Secondly, by employing complete stories as experimental stimuli, this study boosts higher ecological validity than previous well-controlled and sentence-level transmission paradigms, enabling a more comprehensive examination of the dynamic changes in narrative features and brain activity patterns. Moreover, from a data analysis standpoint, this research integrates methodologies from natural language processing and network analysis, allowing for a quantitative representation of various narrative elements such as story content and structure.

However, this study also inevitably has some limitations. Firstly, both the numbers of transmission chains and the generational groups were limited; future research should increase the sample size (number of chains and generations) to enhance the robustness and replicability of the findings, potentially incorporating computational modeling methodologies to explore the information decay, loss, and evolution during the transmission. Additionally, the selection of a single story raises concerns regarding the representativeness of the experimental materials. Future studies should incorporate diverse narrative types to validate whether the structural and content transmission patterns align with the results of this work. Furthermore, excessive head movement during the storytelling phase resulted in the exclusion of data from several participants. Future research should strive to optimize the experimental protocol (e.g., by segmenting the 15-minute experiment into shorter 5-minute intervals, providing participants with sufficient breaks).

In conclusion, this study systematically examined the cognitive and neural foundations of narrative transmission. The findings provide valuable insights into the mechanisms of information propagation, cultural evolution, and the formation of collective memory, offering both theoretical and practical implications.

## Acknowledgments

This research was funded by the National Natural Science Foundation of China (NSFC: 32471100) and Guangdong Basic and Applied Basic Research Foundation (2024A1515030046).

## Author Contributions

**M.Y., L.L., and G.D.** designed the experiment. **M.Y., J.Z., and S.L.** collected the data. **M.Y. and L.L.** analyzed the data, interpreted the results, and drafted the manuscript. **C.L., L.L., and G.D.** provided feedback on data analysis, results interpretation, and manuscript revision. **L.L. and G.D.** supervised the project and edited the manuscript.

## Data and Code Availability

The data and code will be shared upon publication.

## STAR Methods

### Participants

We recruited 58 college students from universities in Beijing. All participants were self-reported right-handed, with an average age of 22.98 ± 2.66 years. To minimize potential gender-related variability in storytelling duration and quality, we included only female participants in this study. To test the stability and reproducibility of the experimental results, two chains of participants were recruited. The first chain, considered the primary analysis, had 39 participants in total, with 9, 10, 8, and 12 participants in each generation, respectively. The second chain, used for testing the reproducibility, had 17 participants in total, with 3, 4, 6, and 4 participants in each generation, respectively. The average word count of the stories narrated by the 58 participants was 952 ± 144 words.

During the phase of listening to stories, two participants were excluded from subsequent MRI data analysis due to excessive head movement (> 3mm or 3 degrees). During the phase of telling stories, seven participants were excluded from the MRI analysis for the same reason. Therefore, during the listening phase, the first transmission chain included four generations with 9, 10, 8, and 12 participants, while the second transmission chain comprised 3, 4, 6, and 4 participants, totaling 56 participants. During the speaking phase, the four generations of the first transmission chain consisted of 9, 7, 7, and 11 participants, and those of the transmission chain consisted of 3, 4, 4, and 4 participants, 49 participants in total.

As this experiment involved tasks related to story comprehension and memory, participants’ story memory ability could be influenced by language proficiency and working memory capacity. Therefore, before the formal experiment, all participants underwent cognitive assessments, including a verbal fluency test (describing pictures) and a working memory test (forward and backward digit span). Before the experiment commenced, all participants provided informed consent, and the study received ethical approval from the Institutional Reviewer Board of Beijing Normal University.

### Experimental Design and Paradigm

In the pre-experiment phase, we selected 7 well-structured and coherent novels. Using a 5-point scale, 27 participants who were not involved in the formal evaluation assessed the familiarity of the stories, the difficulty of memorization and retelling, comprehension complexity, plot intricacy, and the vividness of the stories through an online material evaluation. Finally, we chose stories that had moderate memorizing difficulty and were unfamiliar to the participants as our experimental material. The selected story material was O. Henry’s short story “A Service of Love”, with an average familiarity rating of 1.73 ± 1.13.

In the formal experimental phase, we employed a narrative transmission task that simulated the social transmission process of storytelling. First, one participant (who did not undergo MRI scanning) was recorded when reading the story as the experimental audio material (duration is 5 minutes). Then, the audio was played for the participants in the first generation, who were required to retell the story. Among the first-generation participants, one with higher story retelling quality (close to 5 minutes in duration, with clear audio quality) and better brain imaging quality (head motion within 3mm or 3 degrees, higher signal-to-noise ratio) was randomly selected. After noise reduction processing of this participant’s storytelling recording, her audio was played for the second-generation participants, who were then asked to retell the story. This process was repeated across four generations of participants (see Figure 1A).

In a controlled setting, each participant first listened to a three-minute audio of a classical Chinese poem “静夜思” (“Quiet Night Thought”). This audio was recorded by the same speaker as in the experimental condition, and was played repeatedly for three minutes. Subsequently, participants were instructed to recite the same poem repeatedly within a three-minute time frame. Notably, this classical poem was familiar to all participants from their childhood, allowing them to memorize it effortlessly. The purpose of this control condition was to expose participants to continuous auditory stimuli or to produce words continuously without requiring them to construct new narrative scenarios or consume strenuous memorization efforts. The baseline conditions were part of the dataset but were not used in analyses in the current study.

### Experimental Procedure and Recording

The experimental procedure is depicted in Figure 1A. In the experiment, each participant began with a 30-second resting period, followed by listening to a five-minute narrative. To minimize the influence of non-verbal stimuli, only a headphone icon was presented during the story-listening phase. Subsequently, the screen transitioned to a “+” fixation point, and participants rested for one minute before it changed to a microphone icon, signaling the start of the five-minute narrative retelling session.

Prior to the experiment, participants were informed through instructions that the cue, a flashing microphone icon, would indicate the halfway point and the final 20 seconds of their speaking duration. Participants needed to regulate their narration time and pace based on these visual icon cues. Throughout the experiment, participants were not required to make any button presses or engage in additional tasks.

It was necessary to play and record audio stories during the experiment. Given that MRI machines generate significant noise during operation, which could affect the quality of audio and speech quality and signals for subsequent analysis, this study employed the FOMRI III system (developed by Opto-acoustics, Israel) for audio recording. This system employs machine learning algorithms for online noise reduction in audio signals and effectively reduces background noise from the MRI scanner. To ensure audio quality, after collecting the audio signals within the MRI suite, we further processed the audio files output by FOMRI III using Adobe Audition audio analysis software for offline noise reduction.

### fMRI Parameters and Data Acquisition

The imaging data were acquired by using a Siemens 3T magnetic resonance imaging (MRI) scanner at the Brain Imaging Center of Beijing Normal University. For functional image data acquisition, a T2-weighted gradient echo planar imaging (EPI) sequence was used with the following parameters: TR (Repetition Time) = 2000 ms, TE (Echo Time) = 30 ms, Flip Angle = 90°, FOV (Field of View) = 200 mm, and voxel size = 3.125 × 3.125 × 4 mm.

For structural image data acquisition, an MP-RAGE (Magnetization Prepared Rapid Acquisition Gradient Echo) sequence was employed with the following parameters: TR = 2530 ms, TE = 3.39 ms, Flip Angle = 7°, FOV = 256 mm, voxel size = 1.33 × 1.0 × 1.33 mm.

### Preprocessing

The preprocessing and analysis of the magnetic resonance imaging (MRI) data were carried out using the DPABI toolbox(Yan et al., 2016). The preprocessing pipeline included the following steps: (1) Slice-timing correction, (2) Head motion correction, (3) Co-registration of each participant’s functional images to their individual structural images, followed by normalization to the Montreal Neurological Institute (MNI) space. Functional images were resampled to 3×3×3 mm3. (4) Linear detrending and high-pass filtering at 1/128 Hz were applied. Further denoising included Friston’s 24 head motion parameters (Friston et al., 1996) and five major independent component analysis (ICA) components related to white matter and cerebrospinal fluid signals (Behzadi et al., 2007), followed by Gaussian smoothing with a full width at half maximum (FWHM) of 8 mm.

### Data Analysis

#### Decomposing Narrative Structure and Content

Based on the *Event Segment Theory* (Kurby & Zacks, 2008; Zacks, 2020), a complete story can be divided into several sub-events according to different features (such as places, characters, and plots). Here, the initial version of the story was divided into five different events based on the plots (background, cause, process, outcome, and theme). All the stories told by the participants were also segmented into five corresponding events based on the original story’s segmented events. Both each complete story and each event could be vectorized by using Latent Dirichlet Allocation (LDA) language models. LDA is a common model in topic modeling that characterizes the distribution of latent semantic topics in text, where topics are represented as multinomial distributions over a word vocabulary. Previous research has employed this method for representing text and vocabulary to investigate how the brain represents semantic information (Lyu et al., 2019). As this study used an audio narrative as the experimental stimulus, a novel topic model was adopted for text quantification (Jiang et al., 2018). This model was trained with a large dataset of millions of novels, resulting in a vocabulary size of 243,617 and 500 topics in the LDA model. Therefore, the story text could be represented by model topics, resulting in a 500-dimensional vector representation of the story. This model was trained from Baidu’s open-source project Familia (https://github.com/baidu/Familia).

By computing the cosine distance among event vectors for each story and the relationships and organizational patterns among story events, we transformed complete stories into story networks with the weighted network approach by following previous methods (Lee & Chen, 2022). Individual story events were represented as nodes in the story network, and the edges in the story network represented the correlation and organizational pattern between story events. Consequently, story structure was represented using the edges in the narrative network, while story content was represented by 500-dimensional vectors from the LDA semantic model (Figure 1B). Additionally, the node degree of individual event nodes was used to represent the importance of events within the story. If an event node had higher centrality than other event nodes, it was considered a high centrality event, indicating strong connections and higher semantic similarity with other events. Conversely, a less centralized event was termed a low centrality event. Thus, each complete story’s structural and content representations, along with individual event centralization and semantic characteristics, were vectorized by using LDA language models and network analysis.

For the similarity matrices of structure and content, we computed the correlations among all recalled stories. To assess content similarity, we calculated the cosine similarity between pairs of content LDA vectors, and for structural similarity, we computed the Spearman correlation between pairs of structural vectors. Meanwhile, to evaluate the content and structural fidelity, we calculated the similarity between each participant’s recalled stories and the original story in terms of content and structure, as described above.

To assess whether fidelity varied across generations, we performed an ANCOVA on the fidelity values across the four generations, controlling for differences between chains. Post hoc Bonferroni tests were conducted to evaluate pairwise intergenerational fidelity differences. The same statistical approach was applied to examine variations in within-generation similarity.

Additionally, we utilized a permutation test to compare changing patterns between high- and low-centrality events.

#### Inter-Subject Representational Similarity Analysis

To address the question of how different narrative components are encoded and represented in the brain during the narrative transmission. We employed inter-subject representational similarity analysis (RSA). The IS-RSA method is a model-free method used for identifying brain regions responding to naturalistic stimuli in idiosyncratic ways across various participants (Chen et al., 2020; Finn et al., 2020; Nguyen et al., 2019).

For whole-brain inter-subject RSA, we used a brain parcellation based on meta-analytic co-activations from over 10,000 published studies from (de la Vega et al., 2016), and parcelled each participant’s whole-brain voxels into 50 brain regions. First, for each brain region, we computed the Pearson correlation between the time series of that brain region for all pairs of participants. We then transformed these correlations into dissimilarity values by subtracting them from 1, creating a neural representational dissimilarity matrix (RDM). At the behavioral level, we constructed representational dissimilarity matrices separately for narrative structure and content based on their representation vectors. Finally, to test how much variance of neural dissimilarity matrix could be uniquely explained by content or structure dissimilarity matrices, we constructed a fixed effect model to evaluate the effects of content and structure representations. We repeated the above analysis over 50 whole-brain regions, and all results were corrected for multiple comparisons using false discovery rate (FDR) correction (*q*s < 0.01).

For the following regions of interest (ROI) analysis, we specifically test the neural representations in subareas of the hippocampus and subnetworks of DMN. We parcelled the 400 brain regions from each participant’s whole brain into 17 networks according to (Schaefer et al., 2018) and classified them based on the default subsystems identified by (Andrews-Hanna et al., 2010). This allowed us to extract three subnetworks within the default mode network (Figure 2B and C): the default mode network core system (Core Subsystem, DMNa), the dorsal medial subsystem of the default mode network (Dorsal Medial Subsystem, DMNb), and the medial temporal subsystem (Medial Temporal Subsystem, DMNc). Simultaneously, we divided the hippocampus into anterior and posterior regions (Head and Tail) based on a hippocampal template defined in previous studies (https://neurovault.org/collections/3731/) (Ritchey et al., 2015).

#### Neural Reinstatement and Across-Generation Similarity

Due to these potential functions of the Hippocampus and Default Mode Networks in episodic memory encoding and retrieval (Cohn-Sheehy et al., 2021), we calculated the neural reinstatement within these regions and networks across modalities (listening and speaking), and across generations (participants in the first-generation and left participants) could predict across-generation transmission fidelity.

In terms of neural reinstatement, we computed the Pearson correlation within one brain region within one participant between temporal signals on listening and speaking, and used linear regression models to test if neural reinstatement could predict content and structure fidelity. For the across-generation neural similarity, we first averaged temporal signals across first-generation participants within one brain region and computed the temporal similarity between averaged first-generation signals and participants in other generations. Lastly, we used linear regression models to test if the temporal similarity across generations could successfully predict content and structure fidelity.

#### Functional Connectivity Pattern Similarity

For participants who listened to the original story (first generation), we calculated the functional connectivity (FC) between each participant’s hippocampal subregions and the subnetworks within the default network, performed Fisher’s z transformation, and calculated the average FC pattern as the representation of encoding the original story. Subsequently, we calculated the FC pattern similarity between each participant and the average pattern of the original story (Figure 3A).

We first used fixed-effect linear regression models to estimate the overall effects of FC pattern similarity on predicting the content and structure fidelity, using each FC pattern similarity as a predictor. Then, by removing each FC pattern similarity individually from the model, we further tested the unique variance explained by different FC modes. The significance of each FC mode is illustrated in Figure 3C.

---

**Figure S1.**
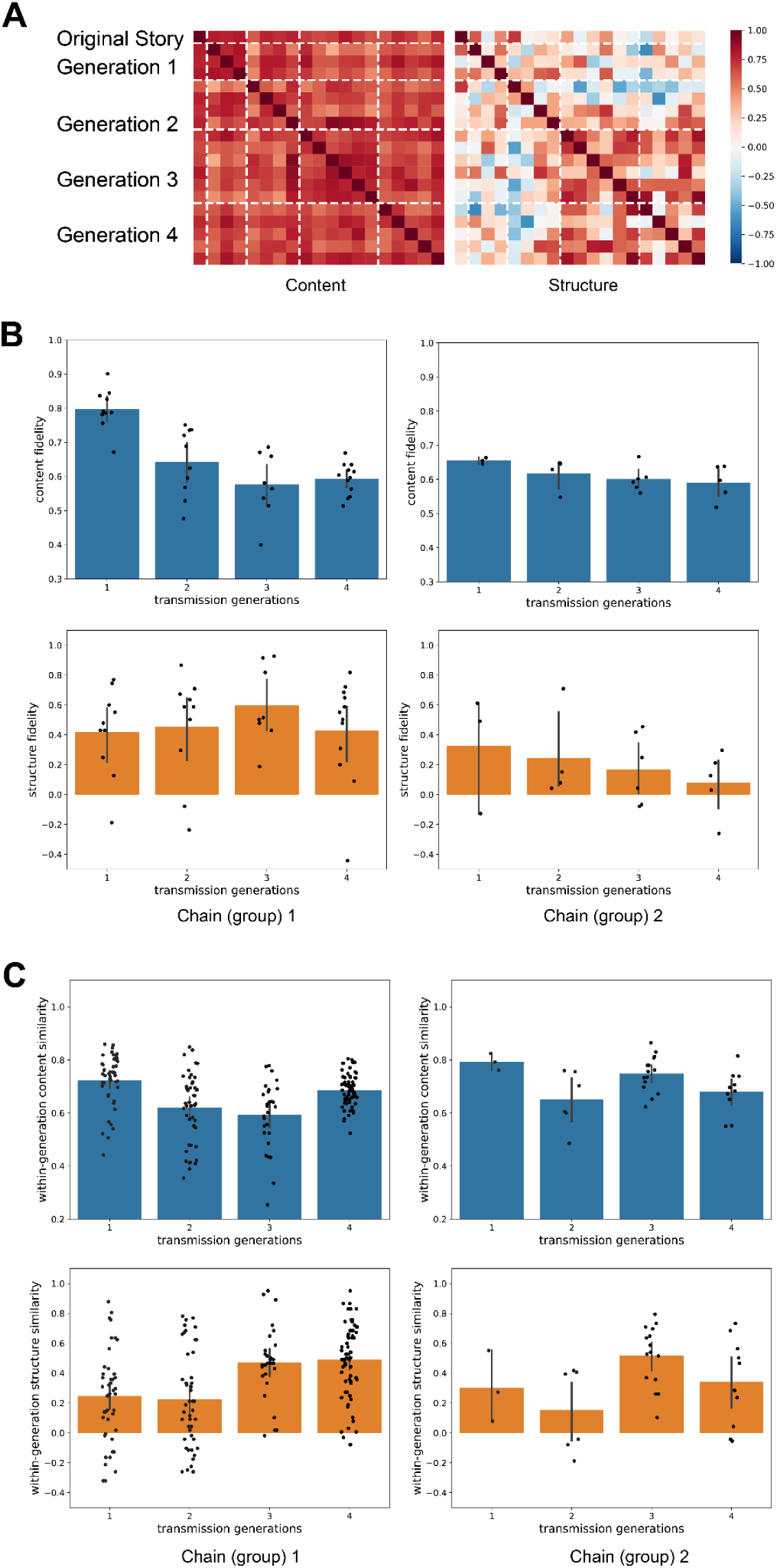
Schematic and Paraphrastic Transmission in Different Chains. (A) Narrative paraphrastic (content, left) and schematic (structure, right) transmission across generations in the second Chain. (B) Chain differences in content and structure fidelity. (C) Chain differences in within-generation content and structure similarity.

**Figure S2.**
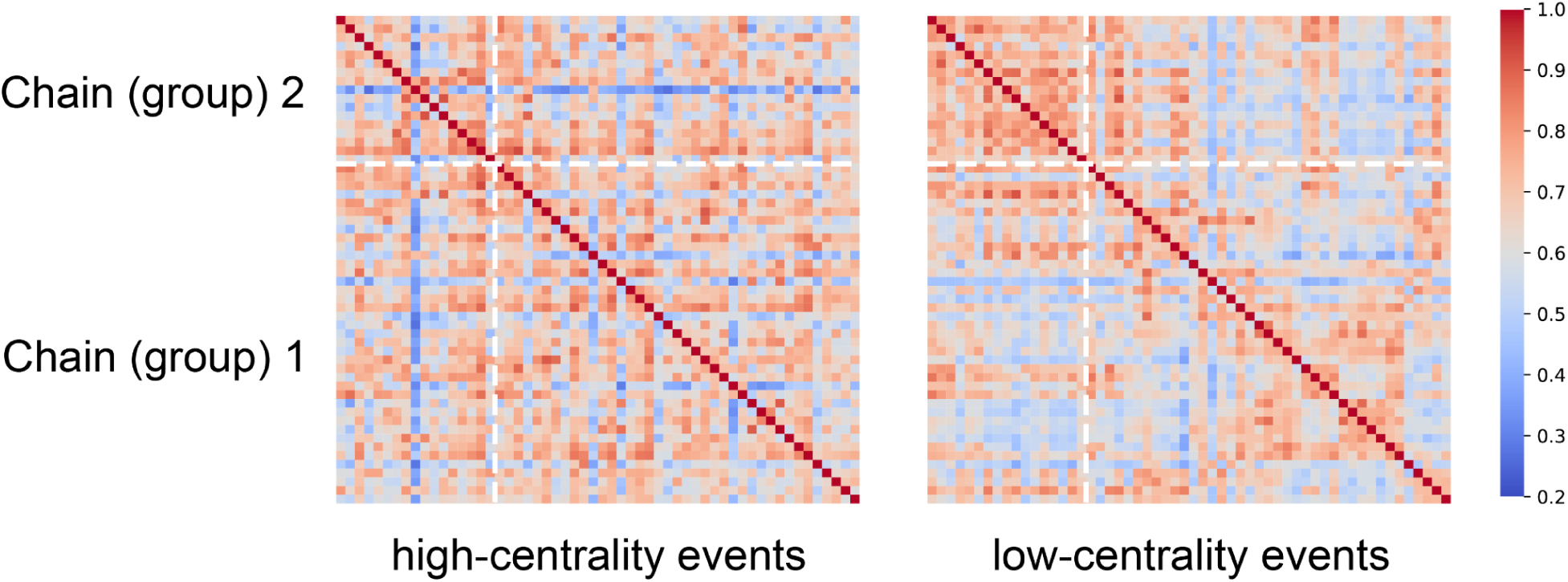
High and Low Centrality Events Transmission. High centrality events showed overall higher similarity during transmission in both chains.

**Figure S3.**
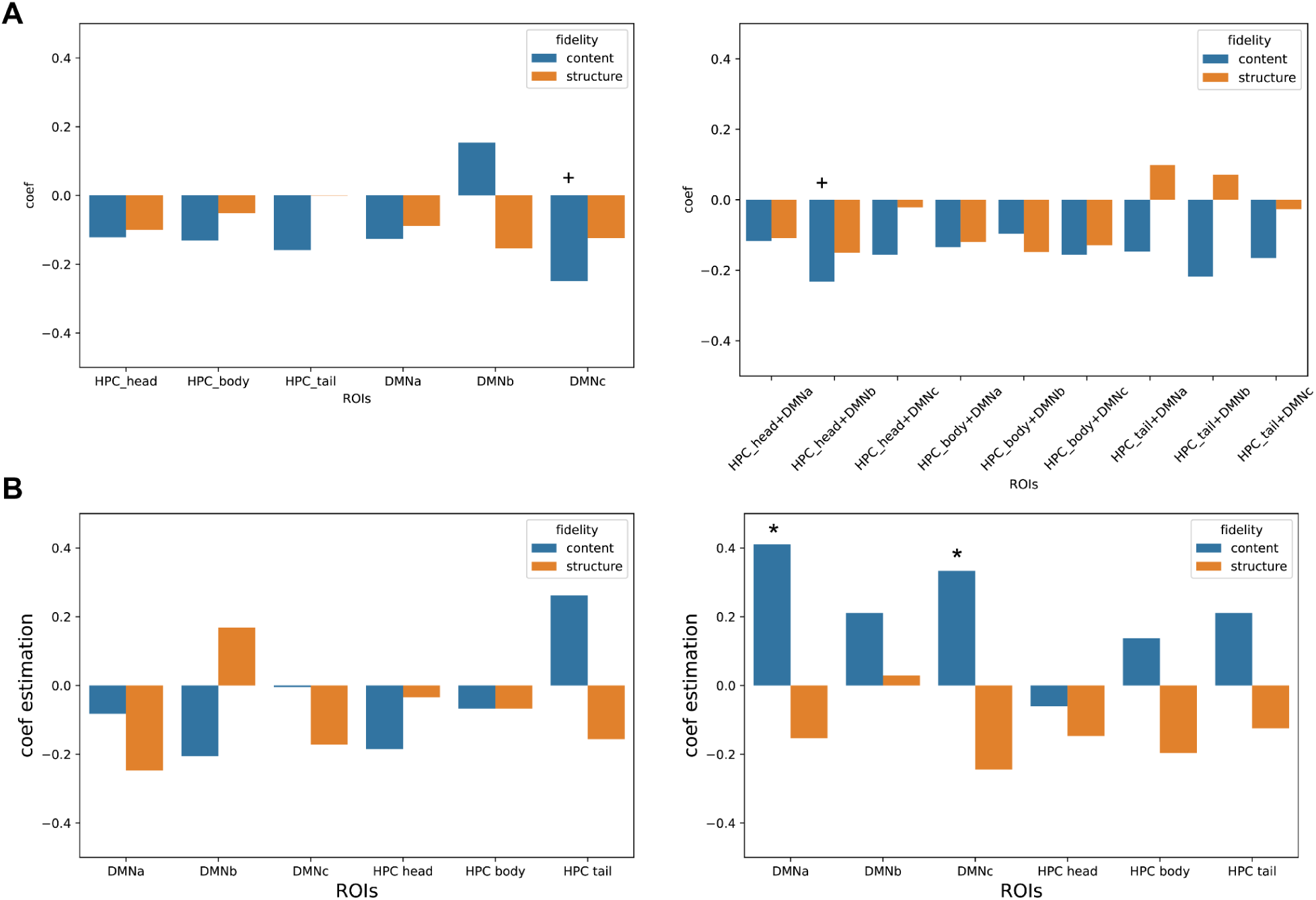
HPC and DMNs Within- and Across-Generation Temporal Similarity Cannot Predict Fidelity. (A) Temporal neural reinstatement between listening and retelling within individuals, either within the same system and region (left) or across different systems and regions (right), showed minimal predictive power for content and structure fidelity. (B) Temporal neural similarity within these ROIs across generations, comparing the average response of the first generation’s to subsequent generations’ responses during listening (left) and speaking (right), could not predict the fidelity of content and structure, despite significant effects of the DMNa and DMNc subnetworks on content fidelity.

## HPC and DMNs Across-Generation Similarity Cannot Predict Fidelity

Previous research indicated that the encoding and retrieval of memories likely occurred through functional connections between the hippocampus and the default network, and this functional connectivity can predict memory performance (Cooper & Ritchey, 2019; Ritchey & Cooper, 2020). Due to these potential functions previous research demonstrated, we further tested (1) if the temporal neural reinstatement between listening and retelling within individuals within or across these systems and regions could predict the fidelity of content and structure; (2) and if the temporal neural similarity within these ROIs across generations could predict the fidelity.

Our results showed that neither temporal neural reinstatement between listening and speaking within one brain region, nor across-generation similarity could significantly predict content and structure fidelity (Figure S3). In detail, we did not find notable evidence that temporal similarity within one subnetwork/ROI between listening and speaking could predict either content (*p*s > 0.277) or structure fidelity (*p*s > 0.291), except for a marginal effect of DMNc on content fidelity (*p* = 0.085) (Figure S3A, left). We further tested if neural reinstatement between listening and speaking across brain regions could predict transmission fidelity (Figure S3A, right). Therefore, correlations between HPC subareas during listening and DMN subnetworks during speaking were calculated within each individual and then utilized to examine their effects on predicting fidelity. Our findings showed that only marginal significance was found on content fidelity and the correlation between HPC head (listening) and DMNb (speaking)(*p*s = 0.099), and apart from this, there is no other significant effect on both content (*p*s > 0.126) and structure fidelity (*p*s > 0.256).

These subtle effects could be caused due to the potential fact that the neural reinstatement might be merely related to the mnemonic effect within one individual and/or one generation, instead of the across-generation fidelity. Therefore, within each brain region/network, we compare the averaged first-generation neural responses (listening to the original story) to the participants in other generations (both listening and speaking), and investigate whether across-generation neural similarity could predict across-generation content and structure fidelity (Figure S3B). We found no significant effects on either content (*p*s > 0.123) or structure (*p*s > 0.146) when comparing averaged first-generation listening neural responses to other participants’ listening signals. Most effects were also insignificant when the averaged first-generation neural signals were compared to other participants’ speaking neural responses, for both content (*p*s > 0.217) and structure fidelity (*p*s > 0.151), other than the significant effects of DMNa and DMNc subnetworks on content fidelity (*p*s < 0.05).

